# Open-Channel Droplet Microfluidic Platform for Passive Generation of Human Sperm Microdroplets

**DOI:** 10.1101/2024.05.09.593416

**Authors:** Tristan M. Nicholson, Jodie C. Tokihiro, Wan-chen Tu, Jian Wei Khor, Ulri N. Lee, Erwin Berthier, John K. Amory, Thomas J. Walsh, Charles H. Muller, Ashleigh B. Theberge

**Affiliations:** Department of Urology, School of Medicine, University of Washington Seattle, Washington 98105, United States; Department of Chemistry, University of Washington, Box 351700, Seattle, Washington 98195, United States; Department of Medicine, School of Medicine, University of Washington, Seattle, Washington 98105, United States

## Abstract

Sperm cryopreservation is important for many individuals across the globe. Recent studies show that vitrification is a valuable approach for maintaining sperm quality after freeze-thawing processes and requires sub-microliter to microliter volumes. A major challenge for the adoption of vitrification in fertility laboratories is the ability to pipette small volumes of sample. Here, we present an open droplet generator that leverages open-channel microfluidics to passively generate sub-microliter to microliter volumes of purified human sperm samples and preserves sperm kinematics. We conclude that our platform is compatible with human sperm, an important foundation for future implementation of vitrification in fertility laboratories.

**Table of contents artwork:** 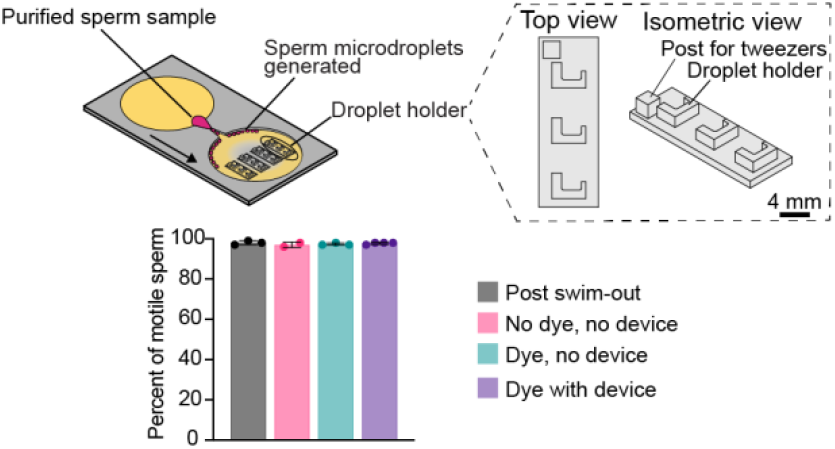

---

Infertility is defined as the inability to conceive after 12 months of unprotected intercourse, or 6 months for couples with a female partner over the age of 35.^1^ Infertility is common, affecting 12.7% of couples in the US or one in six individuals worldwide; a male factor is identified in up to half of couples.^2–5^ Sperm cryopreservation (freezing of sperm for later use) is a component of treatment for many patients with infertility, and also benefits individuals facing military service, individuals planning treatment for cancer, and transgender or nonbinary individuals prior to therapies that impact fertility potential. Traditional sperm cryopreservation techniques (slow and rapid freezing) are associated with adverse impacts on sperm motility, which may decline by 30-50% post-thaw.^6,7^ Thus, there is a clinical need for improvement in sperm cryopreservation to maintain sperm quality. ^8^

Ultra-rapid freezing, or vitrification, is the process of directly inserting small volumes of liquid into liquid nitrogen to reduce ice crystal formation and is the standard approach to cryopreservation of oocytes and embryos.^9,10^ Prior studies have established the benefits of vitrification for sperm preservation^11,12^ and proposed devices that can be used in the vitrification process.^13–15^ A major barrier that has prevented the adoption of vitrification for sperm is the need for generation and manipulation of small fluid volumes (< 1 microliter) to achieve the rapid cooling rates required for vitrification.^16–19^ In standard vitrification workflows, sperm samples are portioned into sub-microliter to microliter volumes using a pipette, which can be time-consuming and challenging to achieve, thus resulting in a need for generating small volumes in an autonomous fashion.^20^

Towards this need, we present a platform to encapsulate purified sperm in microdroplets which leverages hydrostatic and interfacial surface tensions to autonomously generate microliter to nanoliter droplets^21^ without the need for peripheral equipment such as syringe pumps or actuators to drive fluid flow. Our workflow also features a manual step that allows for the manipulation and movement of the droplet through a stylette or forceps into a droplet holder seated in the device, a feature that is only afforded through open channel microfluidics. On a broad scale, microfluidics has been shown to be valuable and fitting for sperm preparation, selection, and analysis for the ability to achieve high-throughput, mimic the female reproductive tract, and enhance assisted reproductive technology (ART) workflows.^22–25^ Open droplet microfluidic devices (including this platform) utilize an air-liquid interface via the removal of the device ceiling, allowing for direct access to the droplets within the device which enables transferring, sorting, moving, and patterning of droplets.^26–31^ In this study, we have generated sub-microliter to microliter sized droplets of purified sperm samples from three healthy sperm donors and demonstrate preservation of sperm motilities and kinematics pre- and post-sample preparation and droplet generation.

The open-platform droplets generator used in this study was developed from Khor *et al*. ^21^ The device with a 0.2 mm wide constriction region was chosen since the expected droplet volumes were less than or equal to 1 µL (Figure 1). Detailed device design and fabrication procedures are in supplemental material. Carrier fluids were prepared from pure HFE-7500 engineered fluid (The 3M Co., St. Paul, Minnesota) with 2 wt% FS-008 fluorosurfactant (RAN Biotechnologies Inc., Beverly, Massachusetts). HEPES buffered formulation with 0.5% Human Serum Albumin sperm wash (#2003, InVitro Care Inc., Frederick, Maryland) was used as the aqueous phase, which was tinted with red dye (McCormick & Co., Hunt Valley, Maryland) at a concentration of 1:500 (v/v) for visualization of the droplets in the device.

**Figure 1.**
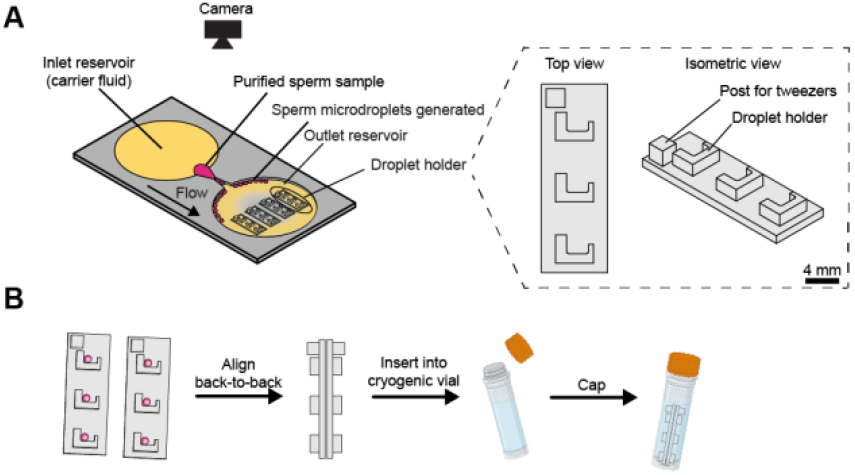
Schematic of open-channel droplet generator and insertion of droplet holder into a cryovial. This device (A) enables the encapsulation of human sperm in microdroplets and the movement from the device to a droplet holder. This holder contains three slots that can hold one droplet each and, after vitrification (B) when placed back-to-back, the holder is able to be placed into a commercial cryogenic vial for long term storage. Created with BioRender.com.

Four droplet holders were then inserted into the slots in the outlet reservoir. A 180 µL aliquot of the dyed sperm wash was added to the inlet platform of the constriction region. Afterwards, 1.5 mL of carrier fluid was pipetted to the larger inlet portion of the device. Once droplet generation began, 0.5 mL of carrier fluid was added to maintain hydrostatic pressure. Devices were then observed for passive droplet generation. Video in supplemental materials shows passive droplet formation.

A stylette or Polytetrafluoroethylene (PTFE)-coated straight-tip forceps (#2A-SA-SE-TC15, Excelta One Star, Buellton, CA) were used to move the droplets from the device to the holder. A single droplet was transferred to each of the droplet holder slots. The droplet holders are designed to be placed back-to-back in a commercial 2 mL polypropylene cryogenic vial (#03374059, Corning Inc., Corning, NY) for long term storage. A video of droplet transfer can be found in the supplementary materials.

Under an approved Institutional Review Board (STUDY00009510) protocol, healthy subjects provided informed consent and semen samples via masturbation following 2-5 days of abstinence.^20^ The semen samples were purified using a direct swim-out procedure.^20^ Motile sperm preparation was added to the device according to Khor *et al*. ^21^ and sperm kinematics were measured for different conditions (with and without dye and passage through the device) using the Hamilton Thorne Research (Beverly, MA) Integrated Visual Optical System (IVOS). A detailed protocol for the swim-out and the device operation can be found in Supporting Information section SI.1.

We have adapted an established droplet generator^21^ (Figure 1A) to be able to seat removable droplet holders (Figure 1A) for the storage of sperm microdroplets in commercial cryogenic vials (Figure 1B). For ultra-rapid freezing methods such as vitrification via liquid nitrogen, sperm specimens which can contain millions of sperm per milliliter require the sample to be fractioned into small volumes (∼1 µL) using a pipette; however, pipetting can be arduous and challenging at low volumes. Our goal for this study is to be able to implement sperm in our device at concentrations that are normal for a healthy patient; encapsulation of sperm at very low concentrations or even a single sperm cell in a microdroplet is not the focus of this work and would only apply to a small percentage of men (1%)^32^. Thus, we leveraged open-channel droplet microfluidics to aliquot purified sperm samples from healthy males with normal sperm counts into microdroplets. The workflow for device operation can be found in Figure 2. We used purified motile sperm samples at a concentration of approximately 6.1 to 7.7 × 10^6^ sperm/mL. A microscope image of a single droplet of the purified sperm can be found in Figure SI.2.1. We have experimentally determined that the mean volume (± standard deviation) of our droplets is 0.8 µL (± 0.3 µL) (Figure SI.3.1). Given a 0.8 µL droplet size, we expect 4.9 to 6.2 × 10^3^ sperm per droplet, which is more than sufficient for *in vitro* fertilization (IVF). As sperm requirements vary among fertility treatments, we can meet different needs through droplet pooling to increase sperm numbers or by adjusting the initial concentration of purified sperm that is loaded into the platform. While our droplets are not monodisperse as have been achieved in closed-channel microfluidics^33–35^, our goal is not to achieve the same degree of monodispersity, but rather to generate small droplets in a passive open system that allows for subsequent manipulation and movement of the droplets. The droplet volumes and dispersity generated by our system are fully acceptable for vitrification for fertility applications.^36^

**Figure 2.**
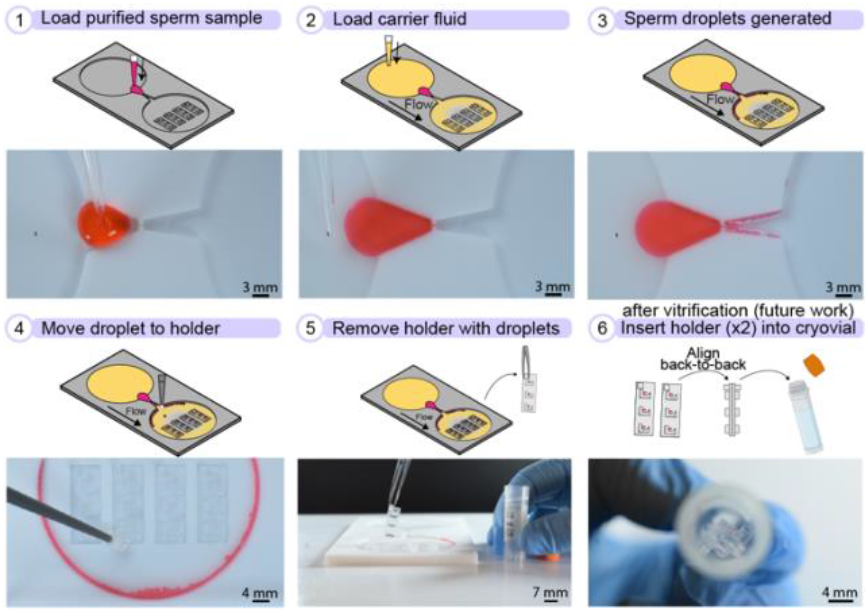
Operation of open-channel droplet generator. A 180-µL sample of purified sperm is loaded with a pipette into the inlet region and carrier fluid consisting of 2% fluorosurfactant in HFE-7500 is subsequently added to the inlet reservoir to initiate droplet generation. Droplets are then autonomously generated, which can be moved to the droplet holder using a stylette or PTFE-coated forceps. Created with BioRender.com.

Within the device, there are four rectangular wells in the outlet reservoir for the seating of the droplet holders. The droplet holder measures 7 mm wide by 20 mm long and 3.92 mm in height and is fabricated out of polystyrene, which is a known biocompatible material. Each slot has a height of 1.50 mm. Each droplet holder can contain three individual droplets and has a handle to allow the holder to be picked up by forceps for vitrification and back-to-back placement into a cryogenic vial (Figure 1B and SI video). Additional droplet holders can be used to store additional droplets depending on the fertility preservation goals of the patient. We tested device usability with users of varying degrees of experience by determining the time needed to fill all droplet holder slots (Figure 3). “Expert” users had prior experience with droplet movement while the “novice” users had basic lab skills, but did not have any experience with the device or droplet movement. Our users were able to move 12 droplets successfully with each user decreasing in time needed with subsequent trials (Figure 3), demonstrating that even new users can operate our device. Currently, manipulation of microdroplets containing human gametes (oocytes) is performed exclusively by highly trained and experienced embryologists using a micromanipulator microscope. Our usability data supports that sperm microdroplet manipulation with our system could be performed by users with basic lab skills. We also examined droplet stability and imaged droplets over the course of 42 minutes after generation. We found that droplets remained in the partitioned state for the entire length of time. Images of the stable droplets can be found in Figure SI.4.1.

**Figure 3.**
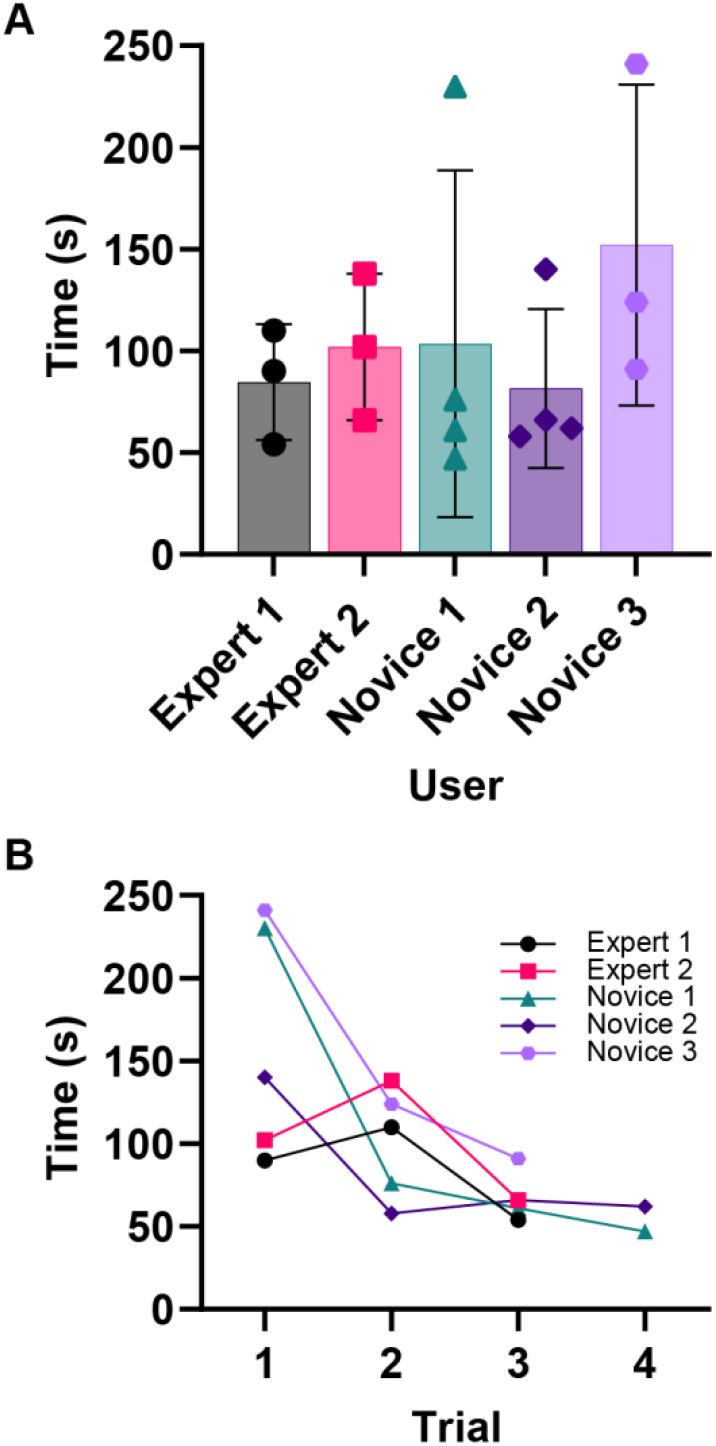
Microdroplets can be successfully manipulated by users of varying experience levels. Users at “expert” and “novice” experience levels were asked to move 12 microdroplets for three or four trials (n=3 or 4). The time needed for each trial was recorded and an average time for each user is shown with the standard deviation (A). (B) A plot showing the progression of each user and the time required to complete the task over each consecutive trial.

**Figure 4.**
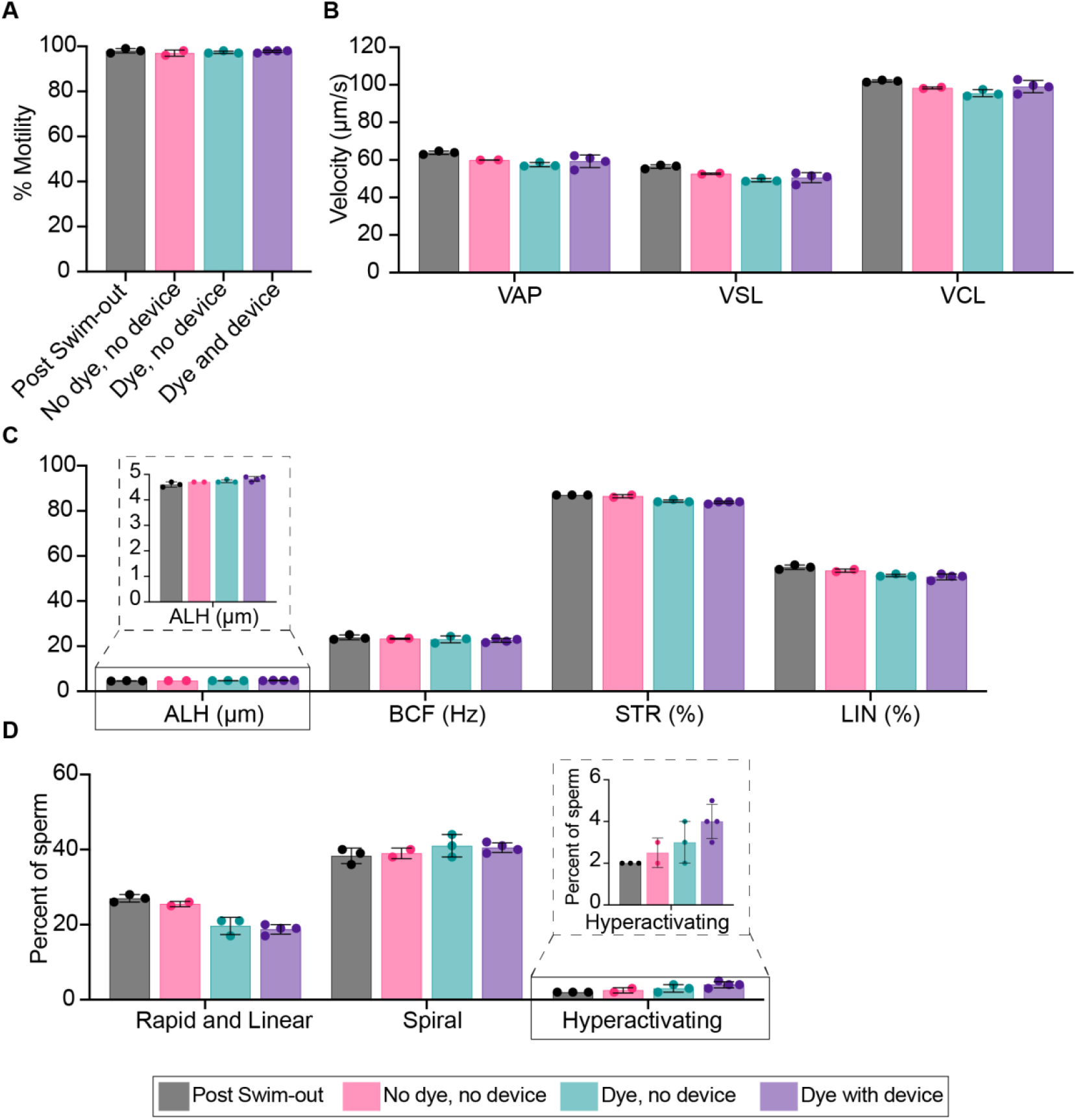
Sperm (A) motilities, (B) velocities, (C) amplitude lateral height (ALH), beat/cross frequency (BCF), straightness (STR) and linearity (LIN) measurements, and (D) movements from one representative participant. The sperm data were collected from post swim-out samples, no dye and no device samples (No dye, no device), with dye (1:500, v/v) without device samples (dye, no device) and with both dye (1:500, v/v) and device samples (dye with device). Three individual participant experiments were conducted with different devices; data from one participant is shown here (Figure 4), and all data is included in the SI (Figure SI.5.1). Each data point represents an individual device and measurement; the bar graph represents the mean ± SD of n = 2-4 devices/measurements.

In this study, the use of dye in sperm samples is crucial for visualization during droplet generation and manipulation. To address this, we investigated sperm motility and kinematic parameters before and after exposure to the dye and the PTFE device. We assessed sperm motility parameters in three sperm donors using an automated sperm analyzer platform, Integrated Visual Optic System (IVOS) (Figure 3 and SI.5.1). A representative participant is presented in Figure 3 and additional participant data can be found in Figure SI.5.1. In these graphs, we present sperm motility data (Figure 3A) as well as additional kinematic measures including average path velocity (VAP), straight-line velocity (VSL), curvilinear velocity (VCL), (Figure 3B) amplitude of lateral head displacement (ALH), beat/cross frequency (BCF), straightness (STR), linearity (LIN), (Figure 3C) as well as percentage of sperm that are rapid and linear, spiraling, and hyperactivating (Figure 3D).^37,38^ We found no notable decline in motility parameters when comparing the sperm samples exposed to dye and the PTFE device or when one or both of those conditions are absent. We have included the post swim-out parameters here, which are measured immediately after swim-out, to account for differences in sperm motilities and kinematics over time, but no apparent differences were found.

Here, we have sustained sperm motility and kinematics using healthy male participants with normal sperm parameters with our adapted open-channel droplet generator. A novel droplet holder feature is also presented, allowing for the storage of droplets in cryogenic vials, which is a highly adaptable feature of our platform. Moreover, our device requires only a pipette, the sperm sample, droplet generation reagents, the device, and droplet holders to operate. With no peripheral equipment necessary and no specialized training needed to generate the droplets, there is strong potential to implement this platform in labs across a broad range of resource levels. Most importantly, our device enables direct access to the droplets for selection and manipulation, which is only afforded through open channels. To our knowledge, our device is one of the few to autonomously generate small volumes for the purposes of sperm vitrification. While our workflow is only autonomous in the droplet generation aspect, we eliminated the need for small-volume pipetting, a technical limitation to sperm preparation for vitrification. Future work may include developing methods to automate droplet manipulation and storage as has been done previously in closed systems.^39,40^ The device and workflow presented here lays the foundation for a new approach to sample preparation for vitrification of sperm, where maintaining sperm motility after passage through the device is crucial in the successful implementation of this device in the clinical setting. Future work will involve performing vitrification of samples generated by our device and comparison of post-thaw sperm quality with conventional methods. Facilitating the adoption of vitrification for sperm could benefit individuals needing fertility preservation, fertility laboratories, and sperm banks.

## ASSOCIATED CONTENT

### Supporting Information

The Supporting Information is available free of charge on the ACS Publications website.

Detailed methods section, image of a droplet with sperm under a microscope, image used in droplet volume calculations, images of droplet stability, and additional participants data (PDF), video of droplet generation (.mp4), and video of droplet transfer (.mp4), design file of droplet generator with holder slots (.STL), design file of droplet holder (.STL)

## Author Contributions

All authors have given approval to the final version of the manuscript.

### Notes

T.M.N., J.W.K., U.N.L., E.B., and A.B.T. filed patent Open Microfluidic Channel Design for Passive Droplet Formation and Manipulation May 23, 2022 through the University of Washington. A.B.T. and U.N.L. report filing multiple patents through the University of Washington and A.B.T. received a gift to support research outside the submitted work from Ionis Pharmaceuticals. E.B. is an inventor on multiple patents filed by Tasso, Inc., the University of Washington, and the University of Wisconsin-Madison. T.M.N. has ownership in Tasso, Inc.; E.B. has ownership in Tasso, Inc., Salus Discovery, LLC, and Seabright, LLC and is employed by Tasso, Inc.; and A.B.T. has ownership in Seabright, LLC; however, this research is not related to these companies. The terms of this arrangement have been reviewed and approved by the University of Washington in accordance with its policies governing outside work and financial conflicts of interest in research. The other authors declare that they have no known competing financial interests or personal relationships that could have appeared to influence the work reported in this paper.

## ACKNOWLEDGMENTS

We thank students and staff of the Male Fertility Lab and the Bioanalytical Chemistry for Medicine and the Environment (BCME) lab for their assistance. Research reported in this publication was supported by the National Institutes of Health National Institute of General Medical Sciences grant R35GM128648 (A.B.T.), National Center For Advancing Translational Sciences grant TL1TR002318 (J.C.T.), KL2TR002317 (T.M.N.), Urology Care Foundation (T.M.N.), the Western Section AUA (T.M.N.) and the University of Washington. We also acknowledge the M.J. Murdock Diagnostics Foundry for Translational Research. The content is solely the responsibility of the authors and does not necessarily represent the official views of the National Institutes of Health.

