## Supporting Information for "Open-Channel Droplet Microfluidic Platform for Passive Generation of Human Sperm Microdroplets"

### Table of Contents


| Section | | Page |
|---|---|---|
| SI.1 | Experimental | S2 |
| SI.2 | Sperm in microdroplet under microscope | S4 |
| SI.3 | Image for droplet volume calculation | S5 |
| SI.4 | Droplet stability images | S6 |
| SI.5 | Participant 1 and 2 sperm kinematic data | S7 |


### Additional supporting information included:

Video: Droplet generation (.mp4)

Video: Droplet holder transfer (.mp4)

Design File: Droplet generator with holder slots (.STL)

Design File: Droplet holder (.STL)

### Section SI.1 – Experimental

#### a. Device design and fabrication

The open-platform generator used in this study was developed from Khor *et al.* [ref]. The device with a 0.2 mm wide constriction region was chosen as the target device since the expected droplet volumes were less than or equal to 1 microliter.

Edits to the open droplet generator and the design of the droplet holder were done in a computer-aided design program (SolidWorks 2023, Waltham, MA). The device design was converted to a computer-aided machining (CAM) “G” code file (.simpl) and post-processed using Autodesk Fusion360 (San Francisco, California). The device and droplet holders were micromilled using a Datron Neo computer-numerical control (CNC) mill. The open droplet generators were milled from 3/16” PTFE sheets (#9266K15, McMaster-Carr, Elmhurst, Illinois). The droplet holders were milled from 4 mm thick polystyrene sheets (#ST31-SH-000200, Goodfellow, Pittsburgh, PA).

Prior to use, the milled devices and droplet holders were cleaned of debris using a soft brush. The parts were then cleaned of any residues by ultrasonication in 70% (v/v) aqueous ethanol for 30 minutes and then sprayed with fresh 70% (v/v) aqueous ethanol. Following the ethanol cleaning steps, the parts were rinsed with deionized (DI) water. Parts were then left to dry in a bioassay dish in the fume hood overnight. Prior to device operation, devices and droplet holders were cleaned of any lint and debris by spraying compressed air for thirty seconds. Devices were then set at a 3-degree incline inside of a bioassay dish using a plastic wedge.

To sterilize the carrier fluid, the solution was syringe filtered using a 0.22 um cellulose acetate syringe filter tip (Whatman, Cytiva, Marlborough, Massachusetts) and a plastic luer-lock syringe (#302995, Becton, Dickson and Company, Franklin Lakes, New Jersey).

#### b. Experimental Observation and analysis of droplet size

Videos of droplet generation were recorded using a Nikon D5300 digital single-lens reflex (DSLR) camera positioned to obtain a top-down view of the device. Videos were recorded at 60 frames per second (FPS).

#### c. Sperm sample preparation and sperm microdroplet generation

Under an approved Institutional Review Board protocol, healthy subjects provided informed consent and semen samples. After 2-5 days of abstinence, samples were obtained by masturbation into a sterile, labeled polypropylene specimen container. All semen samples were incubated at 37°C for 30 minutes prior to analysis. A portion of the semen sample was analyzed for standard semen quality parameters (including concentration and motility) according to World Health Organization protocols (cite WHO Manual of Sperm). All sperm motility assessments were performed using the Hamilton Thorne Research (Beverly, MA) Integrated Visual Optical System (IVOS). A direct swim out procedure was performed by layering 0.5 mL of neat semen underneath 3 mL of Human Tubal Fluid with Human Serum Albumin media (HTF-HSA), and incubation of this preparation for at least 60 minutes at a 30-degree angle at 37°C. The medium containing motile sperm was carefully aspirated and centrifuged at 250 RCF for 7 minutes twice in 3 mL sperm wash media. The final pellet was resuspended in 1-2 mL sperm wash medium and sperm motility was assessed with the IVOS. Experiments with sperm were conducted in the biosafety hood with microchannel placed inside of bioassay dish which was closed when not in use to minimize evaporation. A 1:500 dilution of red food coloring (McCormick) and purified sperm was created and 180 µL of purified motile sperm preparation with 1:500 red food coloring was pipetted into the inlet chamber of the microfluidic channel. 1-2 mL of carrier fluid was pipetted into the inlet and droplet formation was observed. After passage through the microchannel, sperm microdroplets were aspirated and combined into a 7 µL volume for motility analysis with the IVOS. IVOS was also used to assess motility of prepared sperm that were not exposed to red food coloring or the microchannel and red food coloring exposed sperm that were not exposed to the microchannel.

### Section SI.2 – Sperm in microdroplet under microscope

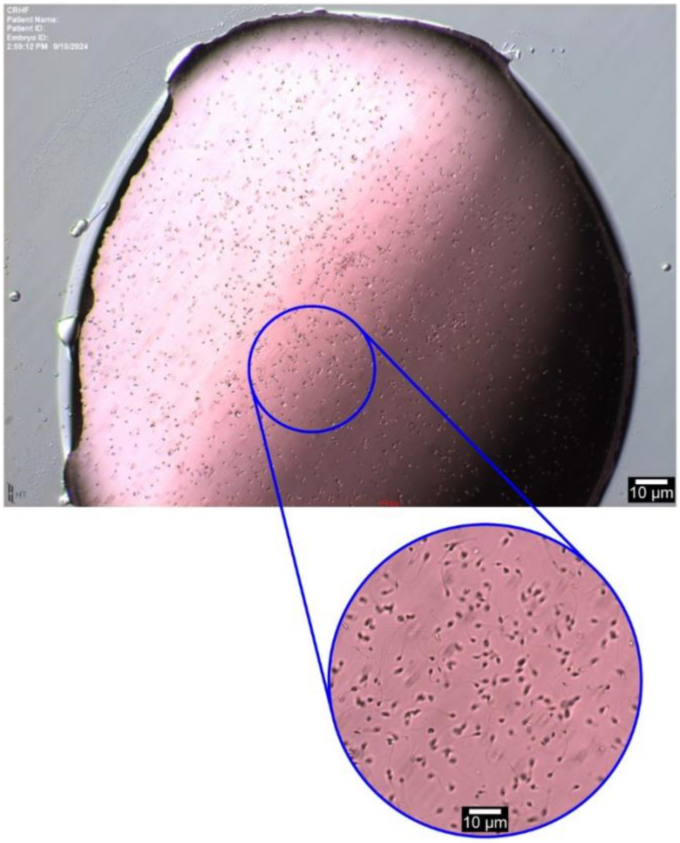


**Figure SI.2.1.** Image of microdroplet containing sperm after droplet generation in the open-channel device taken with a Olympus IX71 inverted phase-contrast microscope (Tokyo, Japan). Microdroplet is dyed red at the same 1:500 concentration (red FC to sperm media) used in the manuscript for visualization and contains approximately 6000 sperm per droplet based on purified sperm sample and median droplet size.

### Section SI.3 – Image for droplet volume calculation

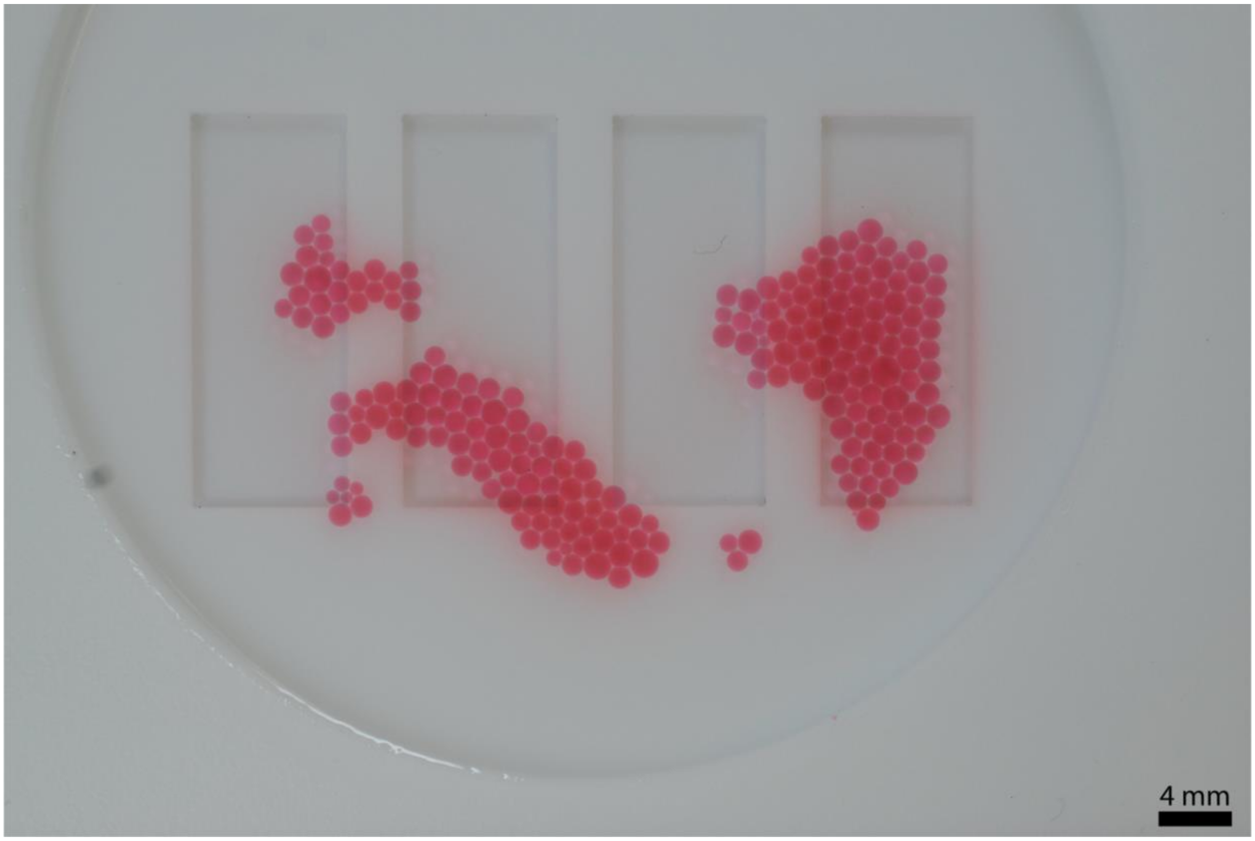


**Figure SI.3.1.** *Droplet volume was calculated using the diameters of the droplets and the equation V = (1/6)(π)(d³).* Droplet diameter was measured using the line tool in ImageJ with three measurements taken per droplet and an average diameter was used for the volume measurement. Volume for each droplet was calculated and averaged to obtain an average volume (± standard deviation) of 0.8 µL (± 0.3 µL). All the droplets generated from one device were used to calculate the average volume.

### Section SI.4 – Droplet stability images

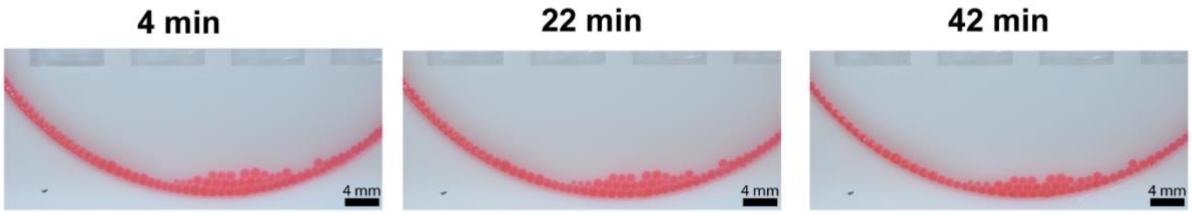


**Figure SI.4.1.** *Droplets are stable for 42 minutes after droplet generation.* Images of dyed sperm media droplets generated at 4 minutes (a), 22 minutes (b), and 42 minutes (c) after the generation of the last droplet. No merging or fusion of droplets was observed during the experiment.

### Section SI.5 – Participant 1 and 2 sperm kinematic data

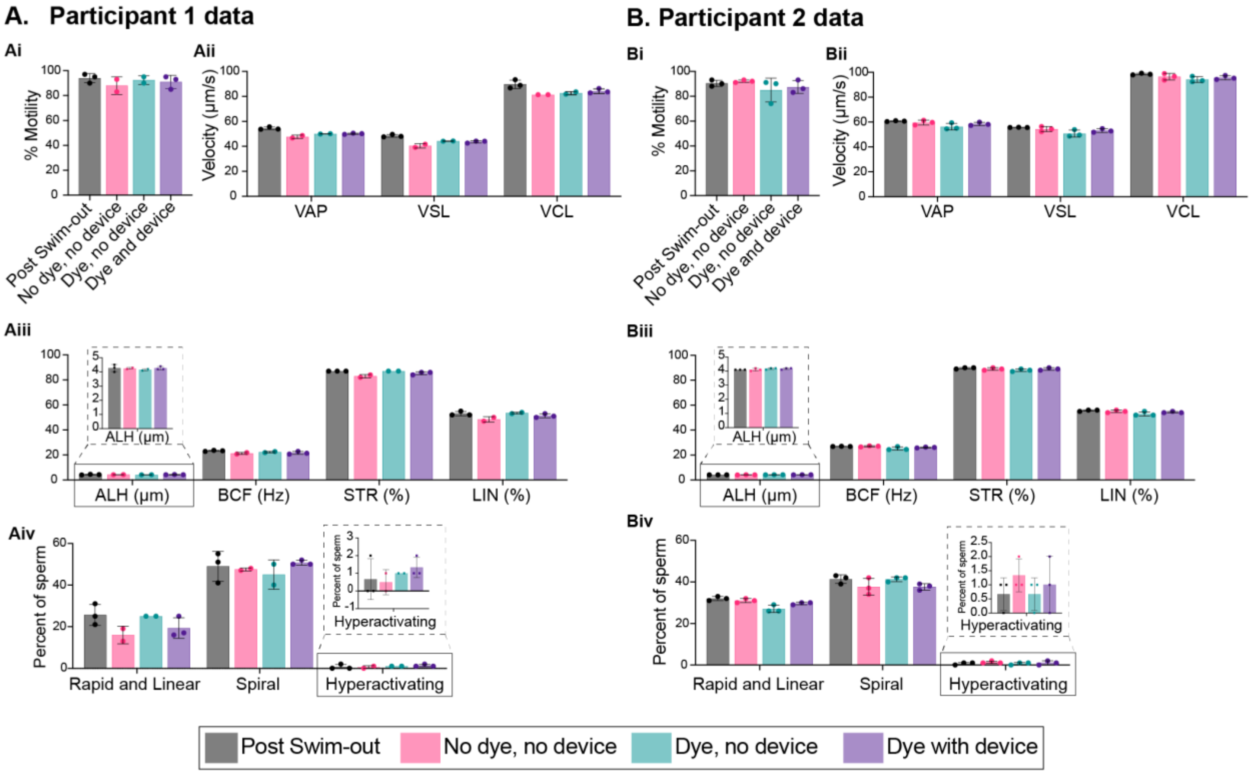


**Figure SI.5.1.** Extended sperm %motility, velocities, and kinematic data were collected with the IVOS. Amplitude lateral height (ALH), beat/cross frequency (BCF), straightness (STR) and linearity (LIN) measurements, and number of rapid and linear, spiral, and hyperactivating sperm were collected for participant 1 (A) and 2 (B). The X axis denotes the measurement with the respective Y-axis units shown in parentheses. Two individual experiments were conducted with the same devices, and the semen samples were from two different participants. Data form a third participant are shown in Figure 4. Each data point represents an individual device; the bar graph represents the mean ± SD of n = 2 or 3 devices.
